# Mobile element warfare via CRISPR and anti-CRISPR in *Pseudomonas aeruginosa*

**DOI:** 10.1101/2020.06.15.151498

**Authors:** Lina M. Leon, Allyson E. Park, Adair L. Borges, Jenny Y. Zhang, Joseph Bondy-Denomy

## Abstract

Bacteria deploy multiple defense mechanisms to prevent the invasion of mobile genetic elements (MGEs). CRISPR-Cas systems use RNA-guided nucleases to target MGEs, which in turn produce anti-CRISPR (Acr) proteins that inactivate Cas protein effectors. The minimal component Type I-C CRISPR-Cas subtype is highly prevalent in bacteria, and yet a lack of a tractable *in vivo* model system has slowed its study, the identification of cognate Acr proteins, and thus our understanding of its true role in nature. Here, we describe MGE-MGE conflict between a mobile *Pseudomonas aeruginosa* Type I-C CRISPR-Cas system always encoded on pKLC102-like conjugative elements, which are large mobile islands, and seven new Type I-C anti-CRISPRs (AcrIF2*, AcrIC3-IC8) encoded by phages, other mobile islands, and transposons. The *P. aeruginosa* Type I-C system possesses a total of 300 non-redundant spacers (from 980 spacers total) across the 42 genomes analyzed, predominantly targeting *P. aeruginosa* phages. Of the seven new Type I-C anti-CRISPRs, all but one are highly acidic, and four have surprisingly broad inhibition activity, blocking multiple distantly related *P. aeruginosa* Type I CRISPR system subtypes (e.g. I-C and I-F, or I-C and I-E), including AcrIF2 (now, AcrIF2*), a previously described DNA mimic. Anti-type I-C activity of AcrIF2* was far more sensitive to mutagenesis of acidic residues in AcrIF2* than anti-type I-F activity, suggesting distinct binding mechanisms for this highly negatively charged protein. Five of the seven Acr proteins block DNA-binding, while the other two act downstream of DNA-binding, likely by preventing Cas3 recruitment or activity. For one such Cas3 inhibitor (AcrIC3), we identify a novel anti-CRISPR evasion strategy: a *cas3-cas8* gene fusion, which also occurs in nature. Collectively, the Type I-C CRISPR spacer diversity and corresponding anti-CRISPR response, all occurring on *Pseudomonas* MGEs, demonstrates an active co-evolutionary battle between parasitic elements.

## INTRODUCTION

The plasticity and rapid evolution of bacterial genomes is driven by the continuous exchange of genetic material between diverse species. This genetic mobility can be blocked by bacterial immune systems, such as restriction enzymes and CRISPR-Cas (Clustered Regularly Interspaced Short Palindromic Repeats and CRISPR associated sequences). CRISPR-Cas systems utilize short RNA guides, encoded within a CRISPR array, where they are separated by repeat sequences, to direct either a multi-protein (Class 1; Type I, Type IIII, Type IV) or single protein (Class 2; Type II, V, or VI) effector complex to a matching target on a mobile genetic element (MGE)^1^. In rare instances, the targeting paradigm is inverted, where a CRISPR-Cas system is encoded by a lytic bacteriophage, targeting the host, as in *Vibrio cholerae*^2^.

*Pseudomonas aeruginosa* is an opportunistic human pathogen and also a leading model organism for studies pertaining to bacteriophage-CRISPR interactions^3^ and Class 1 CRISPR-Cas biology. Functional Type I-F^4,5^, I-E^6^, and now IV-A^7^ systems have been described, however, a fourth CRISPR-Cas system encoded by this species, the Type I-C system has not been well characterized^8^. Type I-C systems are phylogenetically widespread^9^, and can be found in *Streptococcus pyogenes, Vibrio* species, *Clostridium* species, *Neisseria* species, and *Bacillus* species, but are among the least studied subtypes within the adaptive branch of bacterial immunity. Details of Type I-C systems found in *Eggerthella lenta*^10^, *Desulfovibrio vulgaris*^11^, *Bacillus halodurans*^12^, and *Xanthomonas oryzae*^13^ have been explored heterologously or *in vitro*, but studies in a native host are lacking. Type I-C systems employ a minimal surveillance complex of Cas5, Cas7, and Cas8 with the CRISPR RNA (crRNA) and the *trans*-acting nuclease-helicase, Cas3, which is recruited to cleave and processively degrade DNA. The common Cas6 crRNA-processing ribonuclease is missing from this system and Cas5 carries out crRNA-processing instead^14-16^.

Anti-CRISPR proteins (Acrs) encoded by MGEs disable CRISPR-Cas systems using diverse mechanisms. Strategies range from blocking DNA binding sites (e.g. AcrIF1, AcrIF2, AcrIF10, AcrIIA2, AcrIIA4), to blocking DNA cleavage (e.g. AcrIE1, AcrIF3, AcrIIC1) and even acting enzymatically to disable CRISPR-Cas (e.g. AcrVA1, AcrVA5)^3^. CRISPR immunity is typically narrowed to just three stages: adaptation, biogenesis and interference, but a fourth and equally important facet is understanding MGE counter-evolution. Here, we describe the MGE targets of the *P. aeruginosa* type I-C CRISPR-Cas system, which itself is always encoded on an MGE, present direct evidence of endogenous Type I-C CRISPR-Cas activity, and report the discovery of seven *Pseudomonas* Type I-C anti-CRISPRs.

## RESULTS

### MGE-encoded Type I-C CRISPR-Cas provides immunity in *Pseudomonas aeruginosa*

Type I-C CRISPR-Cas systems previously described in 20 *P*. aeruginosa genomes^8^, an environmental isolate in our lab (PaLML1), and 23 additional genomes found using BLAST, are encoded by pKLC102-like elements (Figure 1A). This conjugative element family can be found as either an integrated island or episome in many gram negative bacteria, and is also known as *P. aeruginosa* pathogenicity island (PAPI-1) in some *P. aeruginosa* strains^17,18^. It is typically ∼100 kb, does not always encode a Type I-C system, and we did not observe carriage of other CRISPR-Cas subtypes. To determine if Type I-C CRISPR-Cas is active in *P. aeruginosa*, we first took a bioinformatics approach. While the Cas proteins are conserved (90-100% sequence identity) across strains, the CRISPR spacers are diverse. Alignments of 4,443 protospacers with upstream and downstream regions revealed the consensus PAM to be 3’ –AAG– 5’, consistent with previous reports^10,19^ (Figure 1B). Among the 42 strains with CRISPR arrays published previously (2 published strains have *cas* genes without corresponding arrays), we observed spacer diversity suggestive of active acquisition (Figure 1C and Supplemental Figure 1).

**Figure 1.**
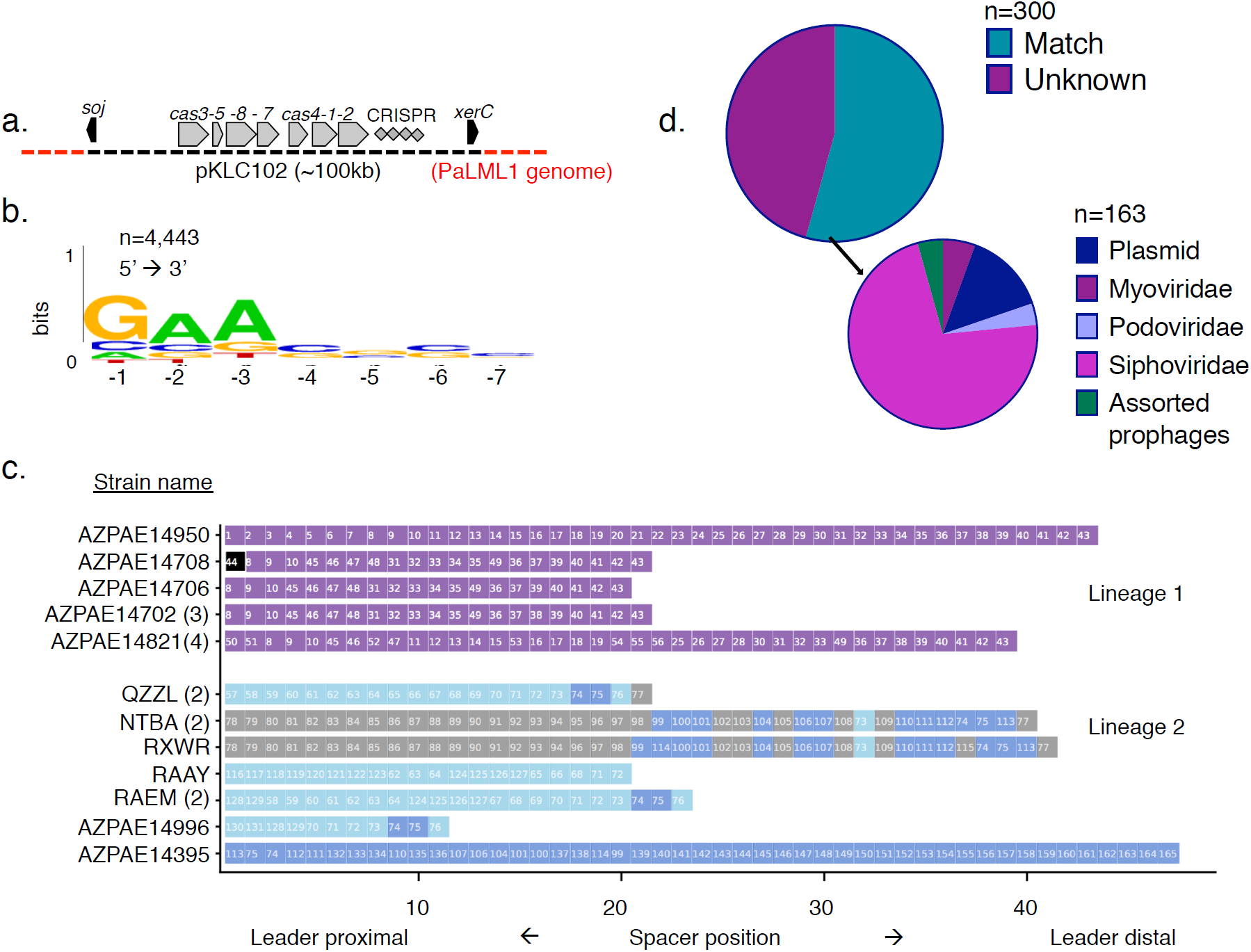
**a**. *Pseudomonas aeruginosa* Type I-C systems are found on pKLC102 elements, shown here integrated into the *P. aeruginosa* genome. Black arrows represent pKLC102 marker genes. *soj* is a chromosome partitioning protein, and *xerC* is a site-specific recombinase. **b**. WebLogo showing the consensus PAM sequence upstream of the protospacer. PAM is written 5’ to 3’. **c**. Clustering of CRISPR arrays from 20 genomes into lineages based on spacer identity. Spacer position is marked on the x-axis. Spacers that are the same within a lineage are given the same number. Numbers in parentheses following the strain names indicate the number of genomes with the same CRISPR array. The spacer highlighted in black, #44, is self-targeting. The colors highlighting the remaining spacers (blue and grey) in lineages 1 and 2 are meant to facilitate comparisons between related arrays. **d**. Of the 300 non-redundant spacers, 163 target sequenced genetic elements. Spacers labeled as unknown (dark purple) did not have any matches in sequence databases used by CRISPR Target. Spacers with matches to independent phage genomes (both lytic and temperate) were categorized into three families (siphoviridae, myoviridae, and podoviridae). Spacers that mapped back to phage-like regions in bacterial genomes were categorized as assorted prophages.

The CRISPR arrays could be clustered into four broad lineages, with strains grouped if they share *at least* one spacer with another array. Strains that cluster together tend to share most of the spacers towards the leader-distal end of the CRISPR array, suggesting that after diverging, each host continues to expand its CRISPR array independently. For example, strains in lineage 1 share most of their ∼10-15 leader-distal spacers, and then undergo obvious divergence with a series of unique spacers proximal to the leader (Figure 1C). In lineage 2, the diversity is even more striking, as the strains are grouped together by just two “core” spacers (#74 and #75), but have highly distinct arrays, most notably strain AZPAE14395, with ∼40 unique spacers (Figure 1C). Strains in lineage 3 (PaLML1, AZPAE14876, and AZPAE12421), and lineage 4 (WH-SGI-V-07071, and WH-SGI-V-07073) have completely dissimilar spacers (Supplemental Figure 1), despite having the same frame shift mutation that results in an early Cas1 stop codon, suggesting continued CRISPR dynamics through an unknown mechanism. In total, there are 300 non-redundant spacers in this collection, and 162 (54 %) match sequenced elements with many spacers targeting phages and prophages (139) and some matching plasmids (23) (Figure 1D). Although pKLC102 can be considered parasitic, dissection of the Type I-C encoded spacers reveals the immunity module to be “domesticated”, targeting canonical bacterial parasites.

A *P. aeruginosa* environmental isolate (PaLML1) from our collection, which has both Type I-C and I-F CRISPR-Cas systems, was next used as a laboratory model. Using WGS data, we determined that PaLML1’s Type I-C system is also within pKLC102 (Figure 1A) and that it clusters with lineage 3, sharing all but one spacer with two of the published CRISPR arrays. To verify CRISPR-Cas function, we transformed PaLML1 with a Type I-C crRNA targeting phage DMS3m, since PaLML1 does not encode spacers against this phage. Because Type I-C spacer length ranges from 32-37 nt, contrary to consistent Type I-F spacers measuring 32 nt (Figure 2A), we tested spacers of each length (i.e. 32 nt, 33 nt, etc.) in PaLML1 to ascertain if all were active. CRISPR-Cas targeting occurred in the presence of the phage-specific crRNAs for both the I-C and I-F systems, and all of the Type I-C spacer lengths tested demonstrated robust phage targeting (Figure 2B). We also isolated escaper phages that had point mutations in positions +2 and +3 of the protospacer (counting from the PAM), suggesting the presence of a “seed” sequence (Figure 2C). In conclusion, active Type I-C systems in *P. aeruginosa* are on a widespread mobile element, have variable CRISPR spacers suggesting activity *in situ*, and can provide protection against phage with an engineered spacer.

**Figure 2:**
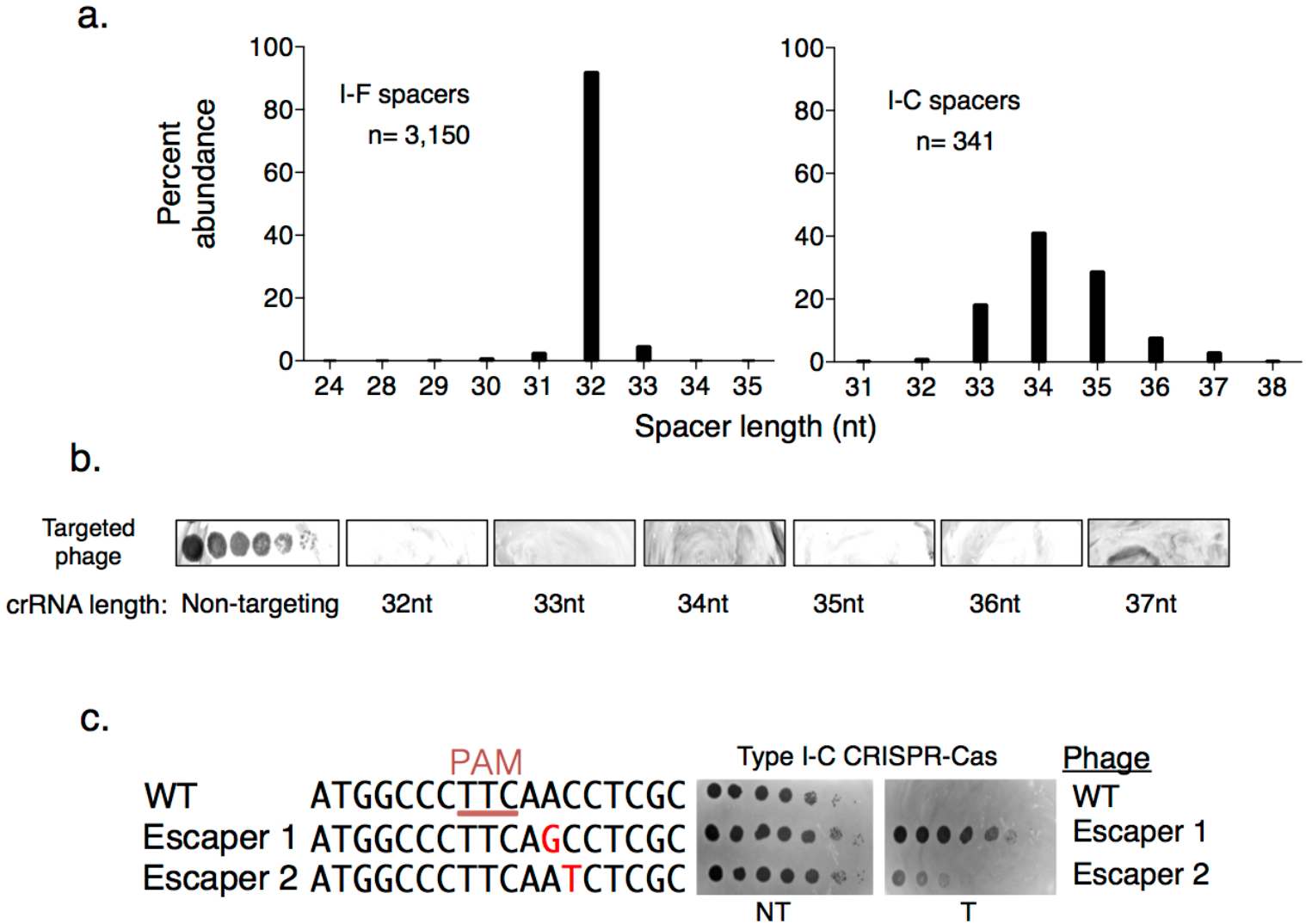
**a**. Comparison of spacer lengths found in either Type I-F or Type I-C *P. aeruginosa* CRISPR arrays. **b**. Spot titration plaque assay of CRISPR-Cas sensitive phage on a lawn of PaLML1, expressing crRNAs of lengths between 32-37 nt. The targeted phage is DMS3m, which does not have an *acrIC* gene. **c**. Protospacer sequence for two Type I-C escaper phages isolated on PaLML1, with mutations highlighted in red text. PAM is underlined. Spot titration plaque assay shows a WT (i.e. non escaper) phage and escapers 1 and 2 challenged with the Type I-C system PAO1^IC^.

### Discovery of seven anti-CRISPRs on MGEs that inhibit Type I-C and beyond

Given the diversity of *P. aeruginosa* Type I-C spacers that target assorted MGEs and the robust phage targeting observed with engineered spacers, we determined that this CRISPR-Cas system indeed poses a threat to MGEs, and therefore counter-immunity mechanisms are expected. To identify candidate anti-CRISPR genes that inhibit this system, we used previously established self-targeting (ST) and guilt-by-association approaches to identify candidates^20,21^. Because cleavage of a bacterial genome is a deadly event^22^, a sequenced strain with a CRISPR-Cas system that has a spacer targeting its own chromosome is indicative of some CRISPR inactivation mechanism allowing that cell to live. Additionally, *acr* genes are often coupled with negative transcriptional regulators known as anti-CRISPR associated (*aca*) genes, which can be used to locate candidate *acr* genes^6,23^. To test candidate Acrs, we used a strain of PAO1 heterologously expressing Cas3-5-8-7 and a DMS3m crRNA from its chromosome (PAO1^IC^)^21^, due to PaLML1’s low transformation efficiency.

Strain AZPAE14708 encodes a spacer targeting its type VI secretion gene, *tagQ*, with a perfect protospacer and PAM match (Figure 3A and Supplemental Figure 2A). This spacer is absent in other strains within lineage 6 that share spacer content with AZPAE14708 (Figure 1B). To identify candidate *acr* genes, we used *acr-*associated gene 1 (*aca1*) as an anchor^6^, and found a locus with *acrIF2*, an inhibitor of Type I-F systems^24^ adjacent to *aca1* (Figure 3A). Surprisingly, expression of AcrIF2 from a phage during infection completely inhibited the Type I-C system (Figure 3B). The dual inhibitory activity was unexpected, given the evolutionary distance between the I-F and I-C systems^1^ (no significant pairwise identity, Supplemental Figure 2B). Two additional AcrIF2 homologues (hereafter, AcrIF2* to indicate dual specificity) were tested (∼50% identity), from *Pseudoxanthomonas* and *Stenotrophonomonas*, both associated with *aca1*, and both displayed dual I-C and I-F activity (Supplemental Figure 2C). Strains from these genera also encode Type I-C and Type I-F systems.

**Figure 3.**
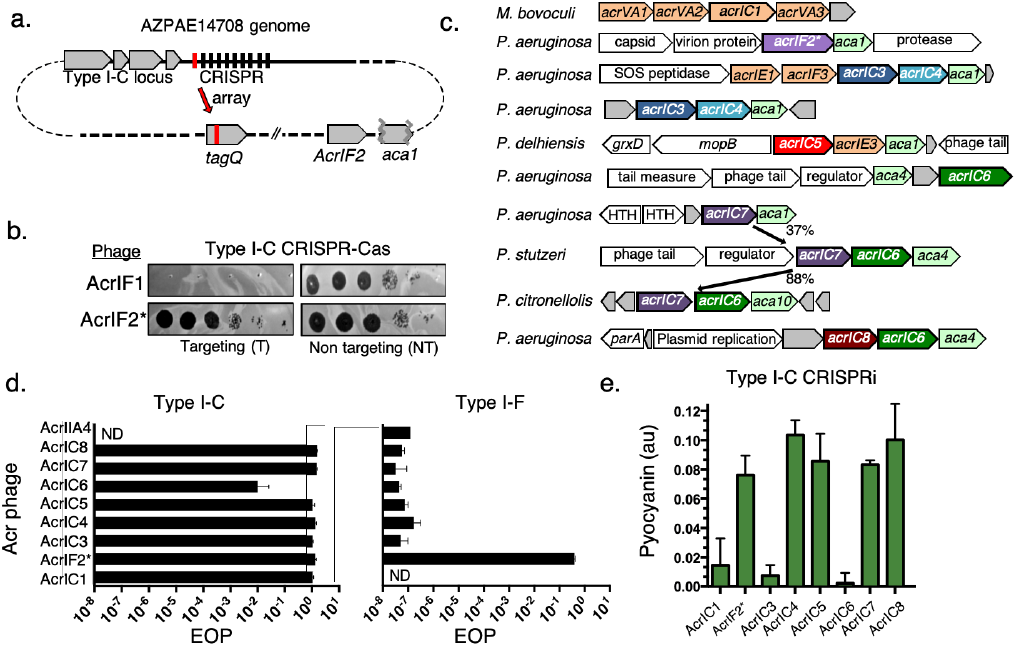
**a**. Schematic of the self-targeting *P. aeruginosa* strain AZPAE14708 showing the first spacer (in red) targeting *tagQ* and the *aca1* locus encoding *acrIF2\**. **b**. A strain expressing the Type I-C CRISPR system in PAO1^IC^ was challenged by phage encoding either AcrIF1 or AcrIF2 in a spot-titration plaque assay with ten-fold serial dilutions. **c**. Gene neighborhood maps of MGEs where new Type I-C *acrs* (colored, bolded arrows) were identified. Previously discovered *acrs* (orange), annotated MGE genes (white), and hypothetical genes (grey), are shown. **d**. Efficiency of plaquing (EOP) calculations for an isogenic panel of phages expressing *acrIC* genes tested in PAO1^IC^ or PA14 (Type I-F). Each strain was infected in triplicate and plaque counts were averaged and normalized against a strain lacking the indicated CRISPR-Cas system. ND-none detected **e**. Transcriptional repression via the Type I-C CRISPR system (CRISPRi, strain: PAO1^IC^ Δ*cas3*) and the impact of the *acrIC* genes. Levels of the pigment pyocyanin are quantified at high levels when CRISPRi is inhibited and low levels when CRISPRi is functional. Each measurement is an average of biological triplicate.

Due to the Type I-C system’s unique mobile lifestyle relative to other CRISPR-Cas systems in *P. aeruginosa*, and AcrIF2*’s narrow distribution, we reasoned that more Type I-C Acrs likely exist. Of 27 *aca1* and *aca4*-assocated candidates tested (Table 1), we identified six more genes in a series of distinct MGEs including plasmids, transposons, conjugative elements, and phages that inhibited the Type I-C system (Figure 3C and Table 2). An additional gene was also identified that solely inhibited the *P. aeruginosa* Type I-E system, AcrIE9 (discussed below). This collection consisted of genes associated with *aca1* (AcrIC3, AcrIC4, AcrIC5) or *aca4* (AcrIC6, AcrIC7, and AcrIC8). AcrIC7 was first identified in *P. stutzeri* (AcrIC7_Pst_) adjacent to *aca4* and a homologue was found in *P. citronellolis* (88% sequence identity, AcrIC7_Pci_), adjacent to a new helix-turn-helix transcriptional regulator, which we have named *aca10*. In both instances, AcrIC6 is also present in the locus. An *aca1*-adjacent distant AcrIC7 homologue was also found in *P. aeruginosa* (37% sequence identity, AcrIC7_Pae_), although it did not confer Type I-C anti-CRISPR activity.

**Table 1:** List of 27 candidates tested, with positive hits highlighted.

| Candidate Number | Accession | Anti-CRISPR identity | aca association |
| --- | --- | --- | --- |
| 1 | KSR23770.1 | AcrIC3 | aca1 |
| 2 | KSO29066.1 | N/A | aca1 |
| 3 | KSL61975.1 | N/A | aca1 |
| 4 | SDK41378.1 | AcrIC5 | aca1 |
| 5 | CDO85538.1 | AcrIC4 | aca1 |
| 6 | WP_085056855.1 | N/A | aca1 |
| 7 | WP_047296680.1 | N/A | aca1 |
| 8 | WP_092238848.1 | N/A | aca1 |
| 9 | WP_044274829.1 | N/A | aca1 |
| 10 | WP_071574229.1 | N/A | aca1 |
| 11 | WP_023657539.1 | N/A | aca1 |
| 12 | ABR13386.1 | N/A | aca4 |
| 13 | ABR13387.1 | N/A | aca4 |
| 14 | SDJ61905.1 | N/A | aca4 |
| 15 | OPE29935.1 | N/A | aca4 |
| 16 | OPD90261.1 | N/A | aca4 |
| 17 | WP_060613673.1 | N/A | aca4 |
| 18 | WP_080050315.1 | AcrIC6* | aca4 |
| 19 | EWC40192.1 | AcrIC7* | aca4 |
| 20 | GCA55691.1 | N/A | aca4 |
| 21 | WP_101192668.1 | AcrIE9 | aca4 |
| 22 | WP_101192667.1 | N/A | aca4 |
| 23 | WP_101192666.1 | N/A | aca4 |
| 24 | WP_045884682.1 | N/A | aca4 |
| 25 | WP_045884679.1 | N/A | aca4 |
| 26 | WP_074202337.1 | AcrIC8* | aca4 |
| 27 | WP_074202338.1 | N/A | aca4 |

\*Multi-subtype Acr proteins.

**Table 2:** Proteins identified and characterized in this study.

| **Acr** | **Size (a.a.)** | **pI** | **CRISPRi phenotype** | **aca association** | **CRISPR-Cas inhibition** | **Accession** |
| --- | --- | --- | --- | --- | --- | --- |
| **AcrIC1** | 190 | 4.17 | Uninhibited | *aca1* | Type I-C | WP_046701304.1 |
| **AcrIF2\*(IC2)** | 96 | 4.02 | Inhibited | *aca1* | Type I-C / Type I-F | WP_015972868.1 |
| **AcrIC3** | 100 | 4.71 | Uninhibited | *aca1* | Type I-C | WP_058130594.1 |
| **AcrIC4** | 57 | 4.22 | Inhibited | *aca1* | Type I-C | WP_153575361.1 |
| **AcrIC5** | 60 | 4.08 | Inhibited | *aca1* | Type I-C | WP_089394111.1 |
| **AcrIC6\*** | 144 | 4.73 | Uninhibited | *aca4* / *aca10* | Type I-C / Type I-E | WP_080050315.1 |
| **AcrIC7<sub>stu</sub>\*** | 94 | 3.85 | Inhibited | *aca4* / *aca10* | Type I-C / Type I-E | WP_003294373.1 |
| **AcrIC8\*** | 80 | 8.01 | Inhibited | *aca4* | Type I-C / Type I-E | WP_074202337.1 |
| **AcrIE9** | 75 | 8.59 | Inhibited | *aca4* | Type I-E | WP_101192668.1 |
| **aca10** | 65 | 8.38 | N/A | N/A | N/A | WP_074980464.1 |

AcrIF2\*(IC2) - AcrIF2 is also the second Type I-C Acr identified, referred to as AcrIF2\* throughout
a.a. – amino acids
\* – indicates dual subtype inhibition.
pI – Average isoelectric point.
CRISPRi – CRISPR interference transcriptional repression assay.
aca – anti-CRISPR associated gene

We subsequently engineered a panel of isogenic DMS3m phages with each individual *acr* gene knocked in, including a negative control (Cas9 anti-CRISPR, *acrIIA4*), regulated by the native DMS3m *acr* promoter and *aca1*, and assessed their efficiency of plaquing in *P. aeruginosa* (Figure 3D). Each phage had an EOP≈1 when infecting cells expressing the Type I-C system, except AcrIC6, which appeared to be quite weak (EOP ≈ 0.01). Only AcrIF2* had activity against the Type I-F system, with an EOP≈1, compared to EOP ≈ 10^−7^ for all other Acr proteins.

To determine how the new Acrs interact with the Cas machinery, we tested whether they inhibit the ability of the crRNA-guided complex to bind DNA *in vivo* using CRISPR transcriptional interference (CRISPRi) in a Δ*cas3* background. A colorimetric assay was adapted from previous work^24^, using a Type I-C crRNA to repress transcription of the *phzM* gene. If CRISPRi is functional, where the surveillance complex assembles and blocks *phzM* transcription, the *P. aeruginosa* culture turns yellow. If DNA-binding is inhibited (CRISPRi negative), the culture is a natural blue-green (Supplemental Figure 2D). Five of the proteins, AcrIF2* IC4, IC5, IC7_Pst_ and IC8, blocked CRISPRi. Expression of AcrIC1 (a previously discovered protein from *Moraxella*^21^) and AcrIC3, however, did not interfere with CRISPRi, suggesting that these Acr proteins bind to Cas3, or prevent Cas3 from cleaving the target DNA, while allowing Cascade-DNA binding (Figure 3E). AcrIC6 did not block CRISPRi but given its weak strength, we are hesitant to interpret this negative result. These results are summarized in Table 2.

### Broad-spectrum inhibitory activity by the I-C anti-CRISPRs

We next surveyed the phylogenetic distribution of the new *acr* genes reported here. AcrIC5 orthologues were found distributed across Proteobacteria, Firmicutes, and Actinobacteria (Figure 4A), and AcrIC8 orthologues were found sparingly in *Pseudomonas*, Spirochetes, and Rhizobiales. AcrIC6 can be found broadly in various classes (Alpha-, Beta- and Gamma-proteobacteria) with notably strong hits in *Salmonella enterica*. These three Acrs stand in contrast to the rest, which were limited to a single genus: AcrIC1 (*Moraxella)*, AcrIC2, AcrIC3, AcrIC4 and AcrIC7 (*Pseudomonas*, data in Table 2). We took note of Actinobacterial AcrIC5 homologues in the human-associated microbes *Cryptobacterium curtum* and *Eggerthella timonensis*, given that an active *Eggerthella lenta* Type I-C CRISPR-Cas system was described recently^10^. We tested the *Pseudomonas* AcrIC5 homologue for inhibitory activity using the established *E. lenta* I-C system heterologously expressed in *P. aeruginosa* and observed strong anti-CRISPR function (Figure 4B), despite *cas* gene sequence identities between 35-55% (Supplemental Figure 3A). Surprisingly, AcrIC7 also inhibited the *E. lenta* I-C system, despite no identified homologues outside of the *Pseudomonas* genus.

**Figure 4:**
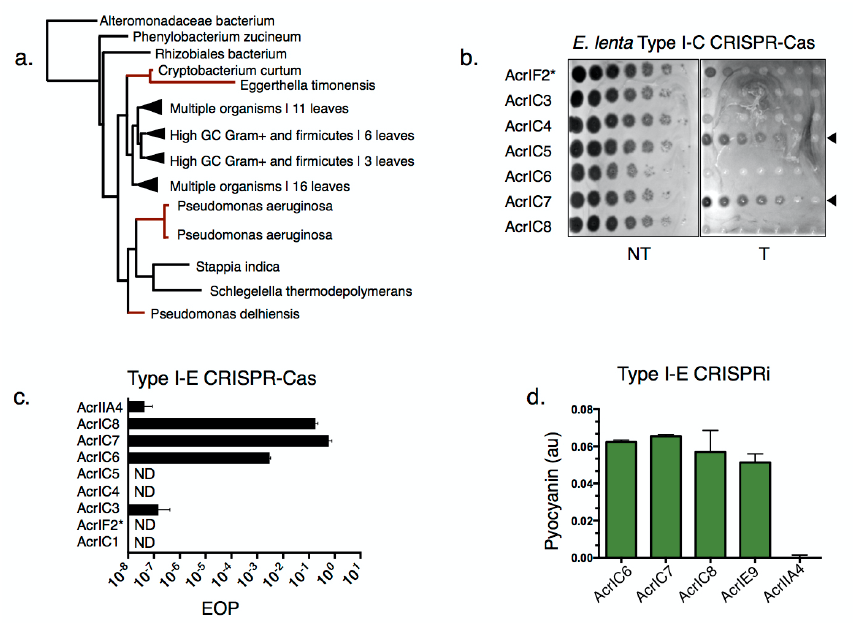
**a**. Phylogenetic tree of AcrIC5 protein showing its broad distribution. **b**. Plaque assay of *acr*-encoding engineered JBD30 phages tested against the *E. lenta* Type I-C system expressed heterologously in *P. aeruginosa*. **c**. EOP calculations for an isogenic panel of phages encoding the indicated *acr* gene, infecting a strain expressing the Type I-E CRISPR-Cas system (PA4386). Each bar is the average of infections done in biological triplicate normalized to the number of plaques on PA4386 Δ*cas3*. **d**. Type I-E CRISPRi, conducted as in Figure 3 (host: PA4386 Δ*cas3*) with the Acr proteins that inhibit Type I-E function assayed. AcrIIA4 is a negative control.

The broad-spectrum activity of AcrIF2 (I-F and I-C), AcrIC5 (I-C_*Pae*_ and I-C_*Ele*_), and AcrIC7 (I-C_*Pae*_ and I-C_*Ele*_), led us to test the new inhibitors against another system found in *P. aeruginosa*, Type I-E. Type I-C, Type I-F, and Type I-E systems are phylogenetically distinct subtypes, with I-F and I-E systems sharing a more recent common ancestor. AcrIC7*_Pst_, AcrIC7*_Pcitro_, AcrIC7*_Pae_, and AcrIC8*, inhibited the Type I-E system well, while AcrIC6* was again, a weak anti-CRISPR (Figure 4C, Supplemental Figure 3B-3D). The new Type I-E Acr proteins (AcrIC6*, AcrIC7*_Pst_, AcrIC8*, and AcrIE9) all inhibited Type I-E CRISPRi (Figure 4D), indicating that they block DNA binding. Curiously, AcrIC7_Pae_ *only* inhibited the I-E subtype, unlike its dual I-C/I-E inhibiting homologues (Supplemental Figure 3C-3E). Searching through sequenced genomes revealed that *P. stutzeri* and *P. aeruginosa* encode both I-C and I-E subtypes, while *P. citronellolis* encodes only Type I-F systems.

### Multi-system inactivation by AcrIF2*

AcrIF2* directly prevents the Type I-F CRISPR surveillance complex from binding to DNA^25,26^. Due to prior structural characterization of AcrIF2*, we opted to next determine whether it uses the same mechanism to inhibit the Type I-C system. Of AcrIF2*’s 96 residues, 24% are acidic, giving it an overall negative charge (pI = 4.0), similar to many of the Acr proteins identified here (Table 2). Despite the Cas proteins from Type I-C and I-F having completely distinct sequences (Supplemental Figure 2B), this negative surface charge could perhaps allow AcrIF2* to block both the I-C and I-F DNA recognition motifs. We therefore conducted structure-guided^25,26^ mutagenesis to attempt to determine whether these two functions could be uncoupled. Eight AcrIF2* residues (D30, E36, D76, E77, E82, E85, E91, E94) were predicted to form key salt bridges between AcrIF2* and Type I-F Cas7/Cas8 (Figure 5A). These were sequentially and incrementally mutated to alanine (starting with a single mutant, then double, and so on), but all of the plasmid-based mutants we tested maintained Acr activity up to an 8xAla mutant (*acrIF2\**^*8xAla*^), while more dramatic mutations (e.g. 8xLys and 8xGlu/Asp) lost function (Supplemental Figure 4). When the 8xAla mutant was expressed from the endogenous phage *acr* locus, we observed that mutagenesis unexpectedly inactivated the anti-Type I-C activity preferentially when infecting PaLML1 (EOP < 10^−4^), while activity against the I-F system was only partially weakened (EOP = 0.02, Figure 5B and 5C). This differential inhibitory activity demonstrates that the mutations impacted one surface-surface interaction more than another, consistent with distinct AcrF2* binding interfaces.

**Figure 5:**
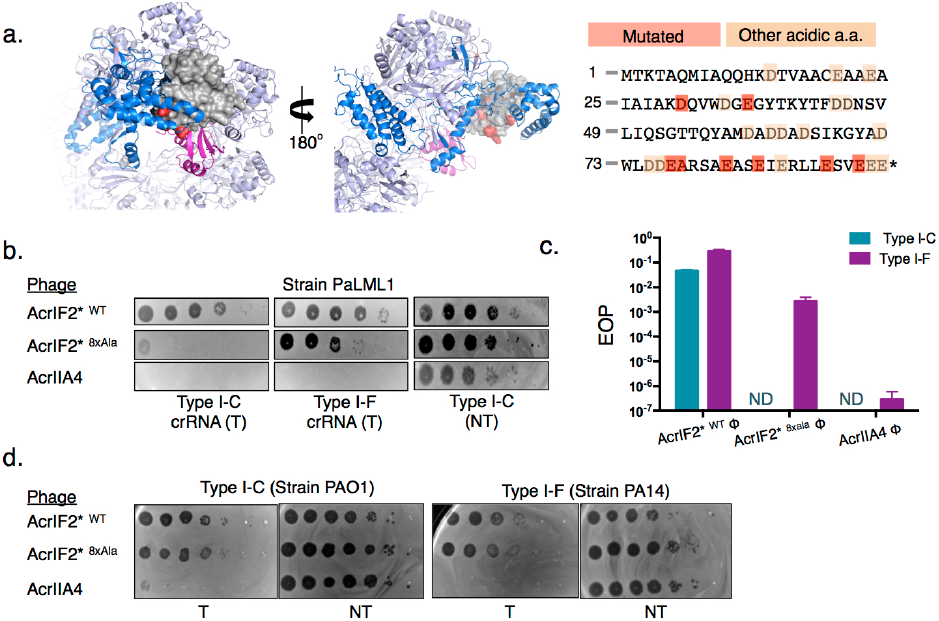
**a**. Color-coded structure of AcrIF2* bound to the Type I-F surveillance complex (PDB: 5UZ9). The Type I-F surveillance complex is shown as a lilac ribbon with Cas8 (blue), and one Cas7 monomer (magenta), AcrIF2* (grey space-filling model), and mutated amino acids (red) shown. AcrIF2* amino acid sequence shown with 8 key acidic residues (red) and all other acidic residues shaded (orange). **b**. Plaque assays with the engineered mutant AcrIF2* phage tested in PaLML1, with either a Type I-F or I-C system crRNA targeting the phage. **c**. Quantification of the efficiency of plaquing (EOP) on PaLML1 for phages expressing the indicated *acr* gene. **d**. Plaque assay with the engineered mutant AcrIF2* phage tested in PAO1^IC^ or PA14. AcrIIA4 is included as a negative control.

Given the dual expression of both I-F and I-C complexes in the PaLML1 strain, we considered whether the weakened activity against the Type I-C system manifests due to weakened binding affinity for that complex coupled with the Acr protein being titrated away by the Type I-F complex. Therefore, we also infected strains that encode just Type I-C (PAO1^IC^) or Type I-F (PA14) with phages encoding WT or mutant *acrIF2\**^*8xAla*^. This revealed that failure of the mutant to inhibit Type I-C function was completely context-dependent as it robustly inhibited the I-C system in PAO1^IC^, which expresses the PaLML1 Type I-C system (Figure 5D). We therefore conclude that while the 8xAla mutant is still capable of Type I-C inhibition, it exhibits a conditional defect in the presence of two competing surveillance complex binding targets in the cell when it’s affinity for the Type I-C system is lowered. These data demonstrate that the highly negative AcrIF2* may use distinct interaction interfaces to enable the inhibition of both the Type I-C and Type I-F CRISPR-Cas systems during infection.

### Anti-CRISPRS that inhibit DNA cleavage by Cas3

Acr proteins that allow for DNA binding but still block phage DNA cleavage, like AcrIC1 and AcrIC3 (Figure 3E), effectively turn the endogenous CRISPR-Cas machinery into a catalytically dead, transcriptional repression (CRISPRi) system. Curiously, AcrIC3 can be frequently found flanked by AcrIE1 and AcrIF3 in *P. aeruginosa*, the only other two Type I anti-CRISPRs that enable CRISPRi^24,27^. This reveals a remarkable “anti-Cas3 locus” for all three Type I CRISPR systems in *P. aeruginosa* (Figure 6A). Conjugative transfer, *parA/B* genes, and type IV secretion system genes are found flanking these *acr* genes. When not found with other CRISPRi-enabling inhibitors, AcrIC3 is carried by phages, along with AcrIC4, which is always paired with AcrIC3.

**Figure 6:**
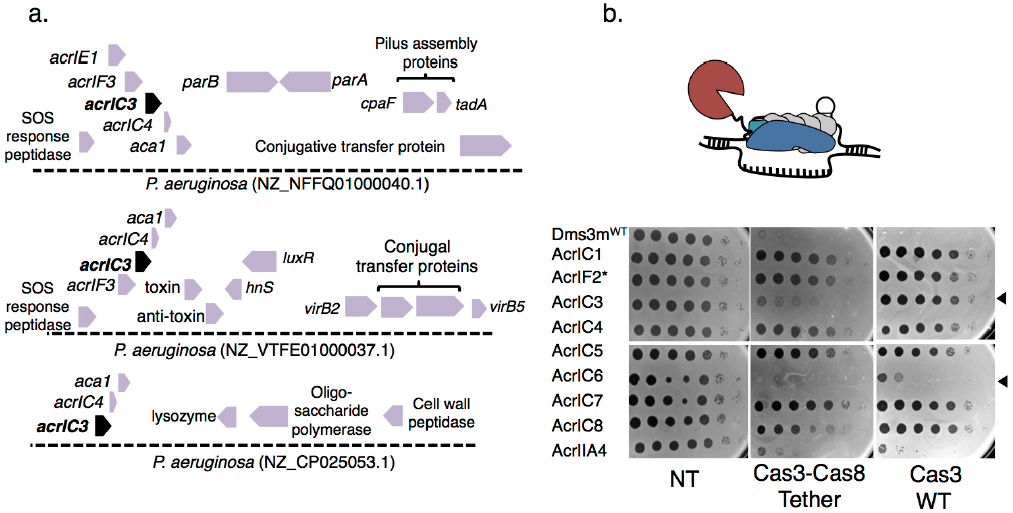
**a**. Gene loci showing *acrIC3. acrIC3* is found on various MGEs, and is often associated with AcrIE1 and AcrIF3, which are Cas3 interacting proteins. **b**. Schematic of the Type I-C mutant where the C-terminus of Cas3 is tethered to the N-terminus of Cas8, with a short linker peptide. Spot titration plaque assay showing the plaquing efficiency of Acr-expressing DMS3m phages on non-targeting (NT), or Type I-C expressing strains, either with Cas3-Cas8 tethered or Cas3 WT.

In an effort to distinguish the inhibitory mechanisms for AcrIC1 and AcrIC3, we constructed a minimal Type I-C complex where the Cas3 C-terminus is tethered to the Cas8 N-terminus with a 13 amino acid sequence (RSTNRAKGLEAVS), effectively granting the surveillance complex nucleolytic activity (Figure 6B). This construct was inspired by, and designed to mimic, naturally occurring variants of Type I-E systems in *Streptomyces griseus*, which encode Cas3 and Cas8 as a single protein, with the same short linker peptide in between^28^. When the panel of Type I-C Acr-expressing phages infected a strain expressing this minimal system, the fusion efficiently evaded the AcrIC3 protein, targeting this phage by ∼1,000-fold, while all other *acr* phages, with the exception of AcrIC6, replicated well (Figure 6B). AcrIC3’s binding site may be occluded with the linker present, or the fusion bypasses a recruitment inhibition mechanism, rendering it an ineffective Acr. Not only does this demonstrate that AcrIC1 and AcrIC3 utilize distinct mechanisms, these data uncover a novel anti-anti-CRISPR strategy in systems with naturally occurring fusions of Cas3 with Cas8.

## Discussion

In the perpetual battle between CRISPR-Cas immunity and genetic parasites, anti-CRISPR proteins are encoded by myriad mobile genetic elements (MGE) to disable CRISPR-Cas activity, allowing for the preservation of the invading element^3^. However, the Type I-C system in *P. aeruginosa* is also mobile, found on a common genomic island (pKLC102) that can exist as either a conjugative island or as a plasmid^8,18,29^. Since mobile elements (here, encoding CRISPR-Cas or anti-CRISPRs) can transfer antibiotic resistance genes, virulence factors, immune systems, and other fitness-altering genetic material to their host^30,31^, this generates an interesting paradigm for CRISPR and anti-CRISPR interactions^32^. These mobile CRISPR-Cas systems can deliver immunity horizontally, granting a recipient a library of spacers against *other* MGEs and the Cas protein machinery. This does not seem to be a rare occurrence, as CRISPR-Cas systems have been identified on plasmids^33^,^34^ and phages^2,35,36^, most notably being used by *V. cholerae* phage ICP1 to neutralize a mobile element with anti-phage activity^2^.

The Acrs described in this study were found encoded by diverse MGEs. AcrIC1, AcrIC2, AcrIC5, AcrIC6*, and AcrIC7* were commonly associated with phage genes, while AcrIC8* is within Tn3 family transposases (Supplemental Figure 3B and 3F). AcrIC3 and AcrIC4 are commonly found together and are associated with D3- and JBD44-like temperate siphophages. AcrIC3 is also common on conjugative elements, where it frequently clusters with Cas3 inhibitors AcrIE1 and AcrIF3. The role of a “anti-Cas3 island” in conjugative transfer from cell to cell is yet to be determined, but this phenomenon may indicate that neutralizing the ssDNAse Cas3 is an effective means to ensure successful conjugative transfer, which proceeds through a ssDNA intermediate.

Of the eight Type I-C anti-CRISPR proteins, all but one (AcrIC8*) had high acidic amino acid content, and are therefore negatively charged at physiological pH (Table 2). This has been a common theme among Acr proteins and inhibitors of other immune systems, which utilize DNA mimicry to block bacterial immunity^37^. Previous AcrIF2* structural work has shown that it partially overlaps with the DNA binding site, thus being considered a DNA mimic or at least a DNA competitor^25,26^. Proteins that mimic DNA can imitate the charge and bend of DNA, which could potentially allow flexibility in inhibiting distinct systems. For example, the T7 phage encoded Ocr protein is highly acidic and forms a dimer with a bend similar to B-DNA^38,39^. Ocr was initially discovered as an effective inhibitor of diverse Type I restriction enzyme systems and was more recently shown to inhibit another anti-phage system, BREX^40^. This suggests that DNA mimicry is a potent and flexible anti-immune strategy. Importantly, systematic mutation of Ocr’s acidic residues revealed it to be highly recalcitrant to breakage, similar to AcrIF2*, maintaining inhibitory activity against Type I R-M even with up to 33% of acidic residues mutated^39^. Similarly, Cas9 inhibitors AcrIIA2 and AcrIIA4 are highly acidic, have broad-spectrum activity^20^, and have been subjected to extensive mutagenesis, also appearing to have dispensable acidic residues^41^. Inhibitor gene over-expression can, however, obscure the interpretation, which is why we placed the *acr* genes under endogenous phage control.

Despite extensive mutagenesis, AcrIF2* retained activity against the Type I-F system and the Type I-C system when each system was expressed separately. While there may be key interactions between an acidic anti-CRISPR and its cognate Cas protein, excess acidic residues could help maintain bonds even when main interactions are broken, and could even hold the key to inhibiting more than one system. Given the robust inhibition of Type I-F in each experiment, charged contacts are perhaps not the main means by which AcrIF2* interacts with the I-F surveillance complex. Hydrogen bonds between AcrIF2* residues proximal to the PAM interacting residues of Cas8 could influence inhibitor activity. If true, AcrIF2* could still be considered a “DNA mimic”, but with different properties than previously suggested. When assayed in a strain expressing both Type I-C and I-F, generating an *in vivo* competition experiment, the 8xAla mutant preferentially lost anti-I-C activity. This suggests that weakened affinity for the Type I-C complex, coupled with >1 unique binding site in the cell revealed a cost to dual-specificity inhibition, at least for the mutant. This result does not conclusively prove that AcrIF2* uses distinct surfaces to disable the Type I-F and Type I-C systems, however, we suspect that this may be the case and await structural analysis of the Type I-C complex and AcrIC proteins.

The role of Acr proteins in the dissemination and maintenance of MGEs in bacterial genomes is just beginning to be explored^42^. Acr proteins facilitate the maintenance of prophages in a genome encoding a spacer against that phage^4,20,43^, which can help CRISPR-Cas be maintained by preventing self-targeting^44^, and even weak Acr proteins can overcome kinetic limitations by working cooperatively^45,46^. The presence of an Acr in a bacterial genome could also confer protection against a CRISPR-Cas system on a mobile element, such as the one encoded by *Vibrio cholerae* phage ICP1^2^, mobile CRISPR-Cas systems on plasmids^7,33^, or islands like pKLC102, where we find the Type I-C system explored in our study. Multi-system inhibition may be a common strategy exploited by MGEs, since bacteria are not limited to only one CRISPR-Cas subtype. Such a tactic conserves genetic real estate, and acts as insurance against the threat of assorted immune systems. Our work underscores the importance of studying CRISPR-Cas vs. Acr mechanisms *in vivo*, and of exploring Acr diversity and mechanisms.

## Acknowledgements

We thank T. Wiegand and B. Wiedenheft (Montana State University) and A. Davidson (University of Toronto) for their thoughtful discussion and feedback. L.M.L was supported by the HHMI Gilliam Fellowship for Advanced Study (HHMI) and the Discovery Fellowship (UCSF). J.B.-D. and the Bondy-Denomy lab was supported by the UCSF Program for Breakthrough Biomedical Research funded in part by the Sandler Foundation, the Searle Fellowship, the Vallee Foundation, the Innovative Genomics Institute, an NIH Director’s Early Independence Award DP5-OD021344, R01GM127489.

## Author Contributions

L.M.L. conducted Acr characterization experiments and spacer and Acr bioinformatics. A.E.P., J.Y.Z., and A.L.B. conducted Acr searches and candidate testing and L.M.L., A.E.P., J.Y.Z., and A.L.B. generated isogenic phage strains. L.M.L and A.E.P. conducted CRISPRi experiments. J.B.-D. conceptualized the project and supervised all bioinformatics and experiments. The manuscript was written by L.M.L. and J.B.-D. with editing and feedback from all authors.

---

**Supplemental Figure 1.**
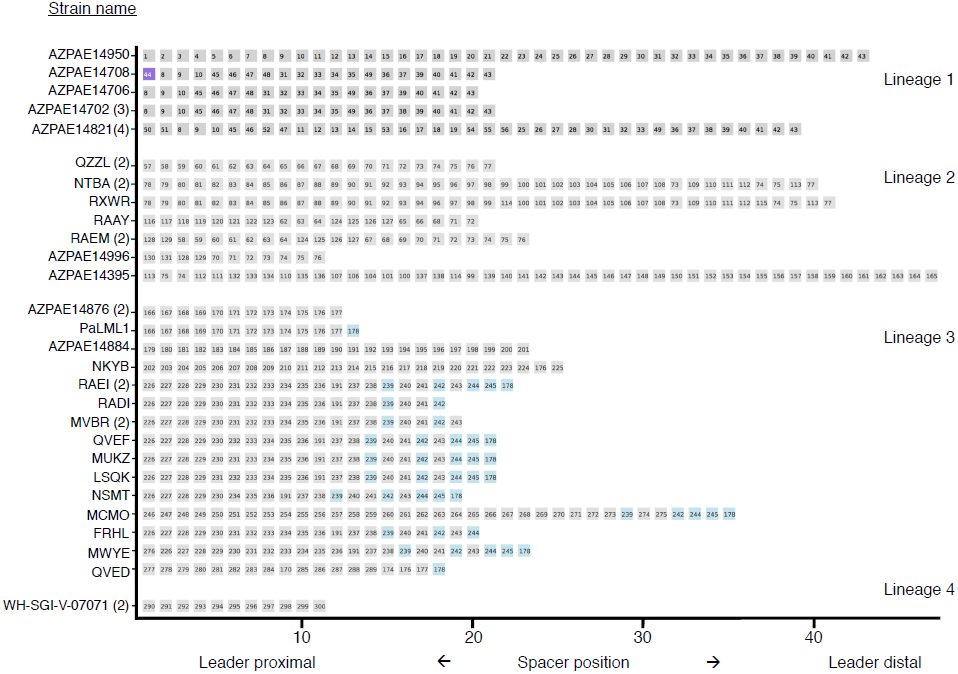
Full CRISPR array lineage mapping of the 28 unique CRISPR arrays from 42 genomes. Each lineage contains CRISPR arrays that share at least one spacer. Spacers with the same DNA sequence are given the same number. Spacer #44 is a self-targeting spacer. Spacers in CRISPR arrays in lineage 3 that are highlighted in blue are meant to facilitate comparisons between related arrays within that lineage.

**Supplemental Figure 2.**
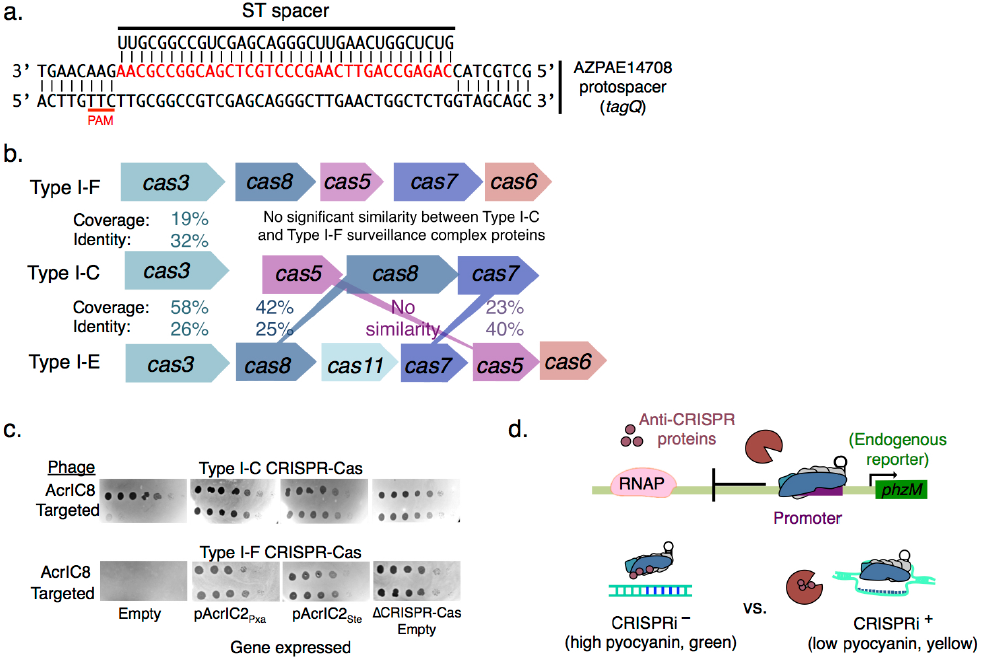
**a**. Alignment of self-targeting spacer #1 from AZPAE14708 with corresponding protospacer. PAM is underlined in red. **b**. Comparison of Type I-F and Type I-E Cas protein sequences to Type I-C Cas protein sequences. **c**. Plaque assay testing the activities of two AcrIF2 homologues identified in *Pseudoxanthomonas* and *Stenotrophomonas* genomes. Homologues were expressed from a plasmid in either a strain encoding the Type I-C system (PAO1^IC^, induced with 1mM IPTG) or the Type I-F system (PA14). A phage encoding a Type I-C Acr (AcrIC8) was used as a positive control, and a phage encoding AcrIIA4 (a Cas9 inhibitor) was used as the targeted phage. **d**. Schematic of the CRISPRi assay used to screen Acr activity. A crRNA is designed to bind upstream of *phzM*, a gene whose expression results in green pigmented *P. aeruginosa* cultures. Acrs that inhibit the surveillance complex from binding target DNA result in a CRISPRi^-^ phenotype. Acrs that bind Cas3 or do not block DNA binding result in a CRISPRi^+^ phenotype.

**Supplemental Figure 3.**
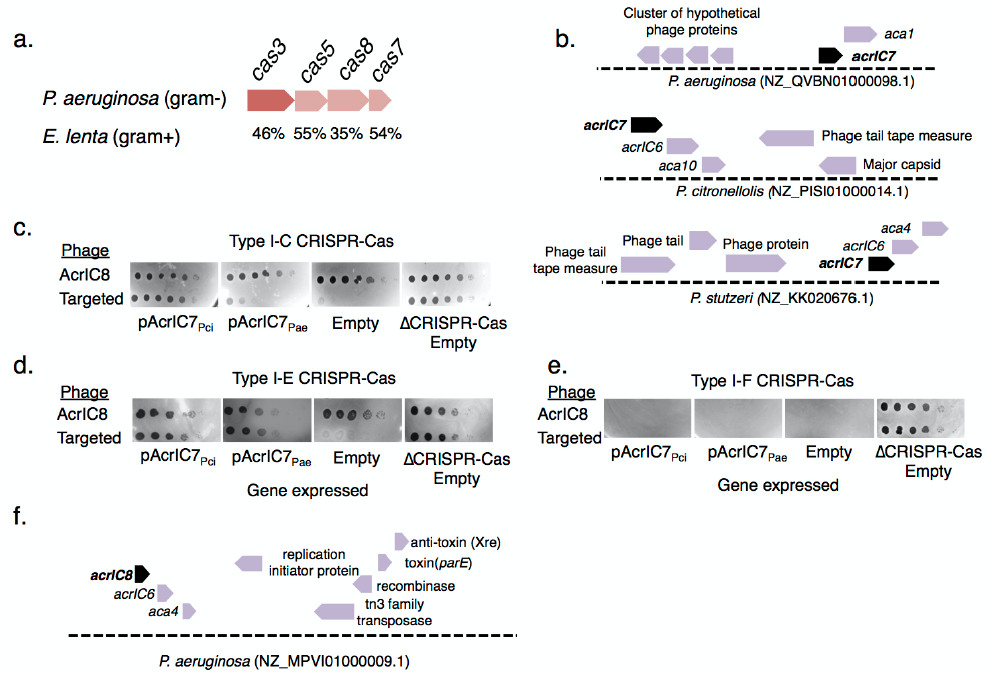
**a**. Protein percent identity comparison of the *E. lenta* Type I-C CRISPR-Cas system to the *P. aeurginosa* Type I-C CRISPR-Cas system. **b**. Loci showing typical genetic context of *acrIC7* in three Pseudomonas species. Genome accession code in parentheses. **c, d, e**. Plaque assays of two AcrIC2 and two AcrIC7 homologues expressed from a plasmid in PAO1^IC^, PA14, or PA4386. Acr activity was assessed by spotting a CRISPR-Cas sensitive phage (DMS3m expressing AcrIIA4) and an untargeted control (DMS3m expressing AcrIC8). **f**. Loci showing typical genetic context of *acrIC8*.

**Supplemental Figure 4.**
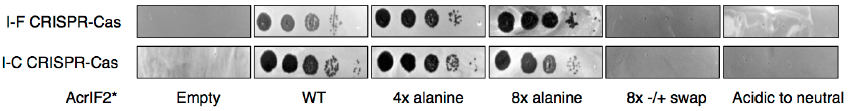
Plaque assays testing the activity of AcrIF2* mutants. A I-F strain (PA14) or I-C strain (PAO1^IC^) were transformed with plasmids encoding the mutants indicated under each panel. A CRISPR-Cas sensitive phage (DMS3m-AcrIIA4) was used to determine the activity of the AcrIF2* mutants.

## Materials and Methods

### Microbes

#### Cell culturing

*Pseudomonas aeruginosa* strains (PAO1, PA14 and PA4386) and *Escherichia coli* strains (DH5a) were cultured using lysogeny broth (LB) agar or liquid media at 37 °C supplemented with gentamicin, where applicable, to maintain pHERD30T (50 µg/mL for *P. aeruginos*a, 30 µg/mL for *E. coli*). In all *P. aeruginosa* experiments, expression of genes of interest in pHERD30T was induced using 0.1% arabinose.

#### Type I-C CRISPR-Cas expression in PAO1

PAO1^IC^ activity was induced using 1mM IPTG. Construction of this strain is described in (*1*) and may be referred to as LL77 (Targeting crRNA) or LL76 (Non targeting).

#### Bacterial transformations

*P. aeruginosa* transformations were performed using standard electroporation protocols (*1*). Briefly, overnight cultures were washed twice in an equal volume of 10% glycerol and the washed pellet was concentrated tenfold in 10% glycerol. These electrocompetent cells were transformed with 20 – 200 ng plasmid, incubated shaking in LB for 1 hr at 37 °C, plated on LB agar with appropriate selection, and incubated overnight at 37 °C. Bacterial transformations for cloning were performed using E. coli DH5α (NEB) according to the manufacturer’s instructions

#### CRISPRi

CRISPR interference transcriptional repression assays were conducted as in previous work (5). A crRNA targeting the *phzM* promoter was introduced into a Δ*cas3* strain. The crRNA and *cas* genes (in the case of Type I-C) were induced in overnight cultures and pyocyanin levels measured with an acid extraction described previously (5).

### Phages

#### Phage maintenance

*Pseudomonas aeruginosa* DMS3m-like phages (including JBD30 and DMS3m engineered phages) were amplified on PA14 ΔCRISPR, PAO1, or PA4386 Δ*cas3* and stored in SM buffer at 4 °C.

#### Construction of recombinant DMS3m acr phages

To generate the isogenic panel of DMS3m and JBD30 anti-CRISPR phages, recombination cassettes were generated with up- and down-stream overhangs to *aca1* and the acr promoter flanking the Acr of interest, as previously described (6). These genes were ordered from TWIST or IDT and were assembled into plasmids using Gibson assembly methods. Recombinant phages were generated by infecting cells transformed with the donor constructs and phages were isolated and assessed for resistance to CRISPR-Cas targeting. The presence of the anti-CRISPR gene was confirmed by PCR.

#### Plaque forming unit quantification

Phage plaque forming units (PFU) were quantified by mixing 10 µl of phage with 150 µl of an overnight bacterial culture. The infected cells were aliquoted into 3 mL molten 0.7 % top agar and spread on an LB agar plate supplemented with 10 mM MgSO4 and appropriate inducers. After 18 hours of growth at 30 °C or 37 °C, individual plaques were counted. Three biological replicates were done per phage per strain.

#### Phage spot assays

3 mL of molten 0.7 % top agar mixed with 150 µl of bacteria were spread on an LB agar plate supplemented with 10 mM MgSO4 to grow a bacterial lawn. Ten-fold serial dilutions of phage were made in SM buffer and 2 µl of each dilution was spotted on the lawn. Plates were incubated at 30 °C or 37 °C for 16 hours and imaged.

#### Efficiency of plaquing (EOP)

EOP was calculated as the ratio of the number of plaque forming units (PFUs) that formed on a targeting strain of bacteria (PAO1^IC^, PA14 WT, PA4386 WT, PaLML1 plus crRNA plasmid) divided by the number of PFUs that formed on a related non-targeting strain (PAO1, PA14 ΔCRISPR, PA4386 ΔCRISPR, PaLML1 plus NT crRNA). Each PFU measurement was performed in biological triplicate. EOP data are displayed as the mean EOP ± standard deviation.

#### Escaper phage isolation

High titer phage preparations were mixed with overnight cultures and spread on an agar plate with top agar. Single plaques that formed after overnight propagation were picked with a sterile pipette tip and resuspended in SM buffer. This process was repeated two times under maintained targeting pressure. The escaper phages were ultimately tittered and the protospacer region sequenced.

### Bioinformatics

Numerical data were analyzed in Excel and plotted in GraphPad Prism 6.0.

#### Discovery of acr genes using *aca1* and *aca4*

Anti-CRISPR searches were done as previously described (*1*)

#### CRISPR array spacer analysis

Spacers were derived from the van Belkum dataset (*2*) (18 genomes with 12 non redundant arrays) or from Type I-C containing strains found using BLAST and CRISPRfinder (*3*)(12 non-redundant arrays). Spacers were analyzed using CRISPRTarget (*4*) using the Genbank-environmental, RefSeq-plasmid, IMG/VR, and PHAST databases.

PAM analysis was done using the Berkeley Web Logo tool by submitting the upstream and downstream regions flanking the protospacer sequence. These 8 nucleotide long flanking sequences are part of the CRISPRTarget output. Every matching protospacer (low cutoff of 20, no redundant matches removed) was utilized for the PAM analysis for n= 4,443.

To determine the types of elements targeted by the spacers in our collection, the cut-off score was increased to 30 and a PAM match score of +5 was used to narrow the total number of hits to matching elements. If a spacer had multiple matches, the match with the highest score was selected as the representative for that spacer. Only one match was considered per spacer. This reduced the number of spacers to 163.

Matches were placed into the following categories: Myophages, Siphophages, Podophages, plasmids, and assorted prophages. A hit was placed into a phage family, rather than into the prophage category, if the CRISPRTarget output included a link to a specific phage genome. Importantly, this means that being placed into a phage family does not mean that a phage is strictly lytic. Prophages were identified by considering the genes in the protospacer neighborhood.

Lineages were manually curated using the 18 strains found in (*2*).

## References

1. Makarova, K. S. et al. Evolutionary classification of CRISPR-Cas systems: a burst of class 2 and derived variants. Nature Reviews Microbiology 18, 67–83 (2020).

2. Seed, K. D., Lazinski, D. W., Calderwood, S. B. & Camilli, A. A bacteriophage encodes its own CRISPR/Cas adaptive response to evade host innate immunity. Nature 494, 489–491 (2013).

3. Davidson, A. R. et al. Anti-CRISPRs: Protein Inhibitors of CRISPR-Cas Systems. Annu. Rev. Biochem 89, 13.1-13.24 (2020).

4. Bondy-Denomy, J., Pawluk, A., Maxwell, K. L. & Davidson, A. R. Bacteriophage genes that inactivate the CRISPR/Cas bacterial immune system. Nature 493, 429–432 (2012).

5. Cady, K. C., Bondy-Denomy, J., Heussler, G. E., Davidson, A. R. & O’Toole, G. A. The CRISPR/Cas Adaptive Immune System of Pseudomonas aeruginosa Mediates Resistance to Naturally Occurring and Engineered Phages. Journal of Bacteriology 194, 5728–5738 (2012).

6. Pawluk, A., Bondy-Denomy, J., Cheung, V. H. W., Maxwell, K. L. & Davidson, A. R. A New Group of Phage Anti-CRISPR Genes Inhibits the Type I-E CRISPR-Cas System of Pseudomonas aeruginosa. mBio 5, e00896 (2014).

7. Crowley, V. M. et al. A Type IV-A CRISPR-Cas System in Pseudomonas aeruginosaMediates RNA-Guided Plasmid Interference In Vivo. The CRISPR Journal 2, 434–440 (2019).

8. van Belkum, A. et al. Phylogenetic Distribution of CRISPR-Cas Systems in Antibiotic-Resistant Pseudomonas aeruginosa. mBio 6, e01796–15 (2015).

9. Makarova, K. S. et al. An updated evolutionary classification of CRISPR– Cas systems. Nature Reviews Microbiology 13, 722–736 (2015).

10. Soto-Perez, P. et al. CRISPR-Cas System of a Prevalent Human Gut Bacterium Reveals Hyper-targeting against Phages in a Human Virome Catalog. Cell Host and Microbe 26, 325–335.e5 (2019).

11. Hochstrasser, M. L., Taylor, D. W., Kornfeld, J. E., Nogales, E. & Doudna, J. A. DNA Targeting by a Minimal CRISPR RNA-Guided Cascade. Molecular Cell 63, 840–851 (2016).

12. Lee, H., Dhingra, Y., Sashital, D. S. The Cas4-Cas1-Cas2 complex mediates precise prespacer processing during CRISPR adaptation. eLife 8, e44248 (2019).

13. Semenova, E., Nagornykh, M., Pyatnitskiy, M., Artamonova, I. I. & Severinov, K. Analysis of CRISPR system function in plant pathogen Xanthomonas oryzae. FEMS Microbiology Letters 296, 110–116 (2009).

14. Nam, K. H. et al. Cas5d Protein Processes Pre-crRNA and Assembles into a Cascade-like Interference Complex in Subtype I-C/Dvulg CRISPR-Cas System. Structure/Folding and Design 20, 1574–1584 (2012).

15. Koo, Y., Ka, D., Kim, E.-J., Suh, N. & Bae, E. Conservation and variability in the structure and function of the Cas5d endoribonuclease in the CRISPR-mediated microbial immune system. J. Mol. Biol. 425, 3799–3810 (2013).

16. Garside, E. L. et al. Cas5d processes pre-crRNA and is a member of a larger family of CRISPR RNA endonucleases. RNA 18, 2020–2028 (2012).

17. Carter, M. Q., Chen, J. & Lory, S. The Pseudomonas aeruginosa Pathogenicity Island PAPI-1 Is Transferred via a Novel Type IV Pilus. Journal of Bacteriology 192, 3249–3258 (2010).

18. Klockgether, J., Wurdemann, D., Reva, O., Wiehlmann, L. & Tummler, B. Diversity of the Abundant pKLC102/PAGI-2 Family of Genomic Islands in Pseudomonas aeruginosa. Journal of Bacteriology 189, 2443–2459 (2007).

19. Leenay, R. T. et al. Identifying and Visualizing Functional PAM Diversity across CRISPR-Cas Systems. Molecular Cell 62, 137–147 (2016).

20. Rauch, B. J. et al. Inhibition of CRISPR-Cas9 with Bacteriophage Proteins. Cell 168, 150–158.e10 (2017).

21. Marino, N. D. et al. Discovery of widespread type I and type V CRISPR-Cas inhibitors. Science 362, 240–242 (2018).

22. Cui, L. & Bikard, D. Consequences of Cas9 cleavage in the chromosome of Escherichia coli. Nucleic Acids Research 44, 4243–4251 (2016).

23. Stanley, S. Y. et al. Anti-CRISPR-Associated Proteins Are Crucial Repressors of Anti-CRISPR Transcription. Cell 178, 1452–1464.e13 (2019).

24. Bondy-Denomy, J. et al. Multiple mechanisms for CRISPR–Cas inhibition by anti-CRISPR proteins. Nature 526, 136–139 (2015).

25. Chowdhury, S. et al. Structure Reveals Mechanisms of Viral Suppressors that Intercept a CRISPR RNA-Guided Surveillance Complex. Cell 169, 47–51.e11 (2017).

26. Guo, T. W. et al. Cryo-EM Structures Reveal Mechanism and Inhibition of DNA Targeting by a CRISPR-Cas Surveillance Complex. Cell 171, 414–419.e12 (2017).

27. Pawluk, A. et al. Disabling a Type I-E CRISPR-Cas Nuclease with a Bacteriophage-Encoded Anti-CRISPR Protein. mBio 8, 43–12 (2017).

28. Westra, E. R. et al. CRISPR Immunity Relies on the Consecutive Binding and Degradation of Negatively Supercoiled Invader DNA by Cascade and Cas3. Molecular Cell 46, 595–605 (2012).

29. Klockgether, J., Reva, O., Larbig, K. & Tummler, B. Sequence Analysis of the Mobile Genome Island pKLC102 of Pseudomonas aeruginosa C. Journal of Bacteriology 186, 518–534 (2003).

30. Thomas, C. M. & Nielsen, K. M. Mechanisms of, and Barriers to, Horizontal Gene Transfer between Bacteria. Nature Reviews Microbiology 3, 711–721 (2005).

31. Oliveira, P. H., Touchon, M., Cury, J. & Rocha, E. P. C. The chromosomal organization of horizontal gene transfer in bacteria. Nature Communications 8, 841 (2017).

32. Faure, G. et al. CRISPR–Cas in mobile genetic elements: counter-defence and beyond. Nature Reviews Microbiology 17, 513–525 (2019).

33. Pinilla-Redondo, R. et al. Type IV CRISPR–Cas systems are highly diverse and involved in competition between plasmids. Nucleic Acids Research 48, 2000–2012 (2019).

34. Crowley, V. M. et al. A Type IV-A CRISPR-Cas System in Pseudomonas aeruginosa mediates RNA-Guided Plasmid Interference In Vivo. The CRISPR Journal 2, 434–440 (2019).

35. Al-Shayeb, B. et al. Clades of huge phages from across Earth’s ecosystems. Nature 578, 425–431 (2020).

36. Medvedeva, S. et al. Virus-borne mini-CRISPR arrays are involved in interviral conflicts. Nature Communications 10, 5204 (2019).

37. Wang, H. C., Chou, C. C., Hsu, K. C., Lee, C. H. & Wang, A. H. J. New paradigm of functional regulation by DNA mimic proteins: Recent updates. IUBMB Life 71, 539–548 (2018).

38. Walkinshaw, M. D. et al. Structure of Ocr from Bacteriophage T7, a Protein that Mimics B-Form DNA. Molecular Cell 9, 187–194 (2002).

39. Roberts, G. A. et al. Exploring the DNA mimicry of the Ocr protein of phage T7. Nucleic Acids Research 40, 8129–8143 (2012).

40. Isaev, A. et al. Phage T7 DNA mimic protein Ocr is a potent inhibitor of BREX defence. Nucleic Acids Research 48, 5397–5406 (2020).

41. Basgall, E. M. et al. Gene drive inhibition by the anti-CRISPR proteins AcrIIA2 and AcrIIA4 in Saccharomyces cerevisiae. Microbiology 164, 464–474 (2018).

42. Mahendra, C. et al. Broad-spectrum anti-CRISPR proteins facilitate horizontal gene transfer. Nature Microbiology 5, 620–629 (2020).

43. Osuna, B. A. et al. Listeria Phages Induce Cas9 Degradation to Protect Lysogenic Genomes. Cell Host and Microbe 28, 1–10 (2020).

44. Rollie, C. et al. Targeting of temperate phages drives loss of type I CRISPR-Cas systems. Nature 578, 149–153 (2020).

45. Borges, A. L. et al. Bacteriophage Cooperation Suppresses CRISPR-Cas3 and Cas9 Immunity. Cell 174, 917–925.e10 (2018).

46. Landsberger, M. et al. Anti-CRISPR Phages Cooperate to Overcome CRISPR-Cas Immunity. Cell 174, 908–916.e12 (2018).

47. Grissa, I., Vergnaud, G. & Pourcel, C. CRISPRFinder: a web tool to identify clustered regularly interspaced short palindromic repeats. Nucleic Acids Research 35, W52–W57 (2007).

48. Biswas, A., Gagnon, J. N., Brouns, S. J. J., Fineran, P. C. & Brown, C. M. CRISPRTarget. RNA Biology 10, 817–827 (2013).

## References

1. N. D. Marino et al., Discovery of widespread type I and type V CRISPR-Cas inhibitors. Science. 362, 240–242 (2018).

2. A. van Belkum et al., Phylogenetic Distribution of CRISPR-Cas Systems in Antibiotic-Resistant Pseudomonas aeruginosa. mBio. 6, 959–13 (2015).

3. I. Grissa, G. Vergnaud, C. Pourcel, CRISPRFinder: a web tool to identify clustered regularly interspaced short palindromic repeats. Nucleic Acids Research. 35, W52–W57 (2007).

4. A. Biswas, J. N. Gagnon, S. J. J. Brouns, P. C. Fineran, C. M. Brown, CRISPRTarget. RNA Biology. 10, 817–827 (2013).

5. Bondy-Denomy, J. et al. Multiple mechanisms for CRISPR–Cas inhibition by anti-CRISPR proteins. Nature 526, 136–139 (2015).

6. Borges, A. L. et al. Bacteriophage Cooperation Suppresses CRISPR-Cas3 and Cas9 Immunity. Cell 174, 917–925.e10 (2018).

